# The G-protein coupled receptor OXER1 is a tissue redox sensor essential for intestinal epithelial barrier integrity

**DOI:** 10.1101/2025.02.05.636712

**Authors:** Miklos Lengyel, Yanan Ma, Zaza Gelashvili, Siyang Peng, Meysoon Quraishi, Philipp Niethammer

## Abstract

Generation of reactive oxygen species is an important part of the innate immune response. Generating microbicidal levels of reactive oxygen species (ROS) requires adaptation of mucosal barriers. High tolerance of ROS provides improved innate immune defenses against pathogens, whereas low tolerance renders host cells prone to chronic toxicity and mutagenesis, which can promote inflammation (e.g., in asthma and Crohn’s disease) and cancerogenesis. The mechanisms that sense and mediate host tolerance to ROS are little understood. In this study, we discover an unexpected role for the redox-sensitive, chemokine-like lipid 5-oxo-eicosatetraenoic acid (5-KETE) in redox adaptation. 5-KETE is known to attract leukocytes to damaged/infected mucosal barriers by signaling through its receptor, OXER1. Suggestive of a distinct non-immune function, we here report that the loss of the OXER1 ortholog Hcar1-4 causes barrier defects and baseline inflammation in the intestine of live zebrafish larvae. In zebrafish and cultured human cells, OXER1 signaling protects against oxidative nucleotide lesions by inducing DNA-protective Nudix hydrolases. Our data reveal the oxoeicosanoid pathway as a conserved ROS resilience mechanism that fortifies pathogen-exposed mucosal linings against increased oxidative stress *in vivo*.

## Introduction

At sites of mucosal damage, large amounts of reactive oxygen species (ROS) are produced to combat invading microbes [1]. Although effectively damaging pathogens, ROS can also harm host cells. To avoid excessive self-injury, the host must thus coordinate ROS production with protective and regenerative mechanisms. Those may include the recruitment of repair leukocytes (N2/M2 neutrophils or macrophages), stimulation of growth factor signalling, and upregulation of antioxidant defenses (e.g., glutathione peroxidases, Nudix hydrolases, etc.) [2, 3]. This coordination involves monitoring metabolic or biochemical indicators of oxidative stress—such as NADP^+^/NADPH ratios, thiol-, and lipid oxidation — through “redox signaling” [4, 5]. Most current research on redox signaling focuses on the posttranslational modification of kinases and phosphatases by reversible thiol oxidation [6]. However, because polyunsaturated fatty acids like arachidonic acid (AA, eicosatetraenoic acid, C20:4) contain bis-allylic methylene groups that are highly prone to both enzymatic and non-enzymatic oxidation, the metabolism and receptor interactions of cell-permeable signaling lipids—such as 5-KETE (5-oxo-eicosatetraenoic acid)—represent another key pathway for reversible redox signaling in tissues. Oxidative stress increases the production of 5-KETE [7–9], and elevated levels of 5-KETE have been found in tissue samples from patients with asthma, chronic obstructive pulmonary disease, or pulmonary hypertension [10–12]. Unraveling 5-KETE/OXER1 biology *in vivo* has been hampered by the lack of OXER1 orthologs in rodents. In contrast, zebrafish express the OXER1 ortholog Hcar1-4 and are a powerful model to study innate immunity and leukocytic inflammation at mucosal lesions. Zebrafish thus had been key to discover OXER1’s conserved physiological role in rapid immune defense at epithelial wounds and infection sites. Most work on the oxoeicosanoid pathway, so far, pertained to its immune functions [13, 14]. Aiming to dissect the immune functions of Hcar1-4 in mucosal inflammation *in vivo* using a previously established zebrafish colitis model [15, 16], we serendipitously discovered an unexpected, phylogenetically conserved function of oxoeicosanoid signaling in the redox protection of mucosal linings.

## Results and Discussion

To study intestinal oxoeicosanoid signaling, we chemically induced mucosal inflammation with dextran sodium sulphate (DSS) in live zebrafish larvae as previously described [15, 16]. DSS increased intestinal neutrophil infiltration in wild-type larvae (Figure 1A, top panels) as expected. Surprisingly, *hcar1-4* deletion larvae (*hcar1-4^mk214/mk214^*) exhibited high neutrophil infiltration at baseline that was not further increased by DSS (Figure 1A, lower panels, also see quantification). A similar phenotype was observed in a different *hcar1-4* mutant line (*hcar1-4^mk213/mk213^*, Supplementary Figure 1A). By contrast, the deletion of 5-lipoxygenase (*alox5a*), an enzyme involved in the production of 5-KETE from arachidonic acid did not affect baseline neutrophils counts, but completely blocked DSS induced neutrophil recruitment (Supplementary Figure 1B). This is consistent with the known pro-inflammatory role of Alox5a after epithelial injury [17]. We conclude that loss of Hcar1-4 causes spontaneous neutrophilic inflammation. Our published RNA sequencing data of *hcar1-4* mutant embryos [13] revealed the increased expression of cell death-related genes (e.g., *casp6a*) in *hcar1-4^mk213/mk213^* larvae. A pronounced increase in intestinal apoptosis was observed in both *hcar1-4* mutants (Figure 1B, Supplementary Figure 1C). Simultaneously, gut barrier leakiness was increased in *hcar1-4^mk214/mk214^* intestines (Figure 1B). The pan-caspase inhibitor ZVAD-FMK normalized the baseline neutrophil numbers in the *hcar1-4*^mk214/mk214^ intestines (Figure 1C). To distinguish whether apoptosis was a result of increased neutrophilic inflammation or vice versa, we chemogenetically ablated nitroreductase (NTR2.0) expressing neutrophils in *hcar1-4* mutant fish by bathing the larvae in metronidazole [18], which did not reduce intestinal apoptosis (Figure 1D). Likewise, broad spectrum antibiotics (Abx) normalized intestinal neutrophil counts in the *hcar1-4* mutant larvae without suppressing apoptosis (Figure 1E). These data suggest that *hcar1-4* deletion causes intestinal apoptosis and leakiness along with opportunistic bacterial infection, and that the increased neutrophilic inflammation at baseline is downstream of chronic infection.

**Figure 1.**
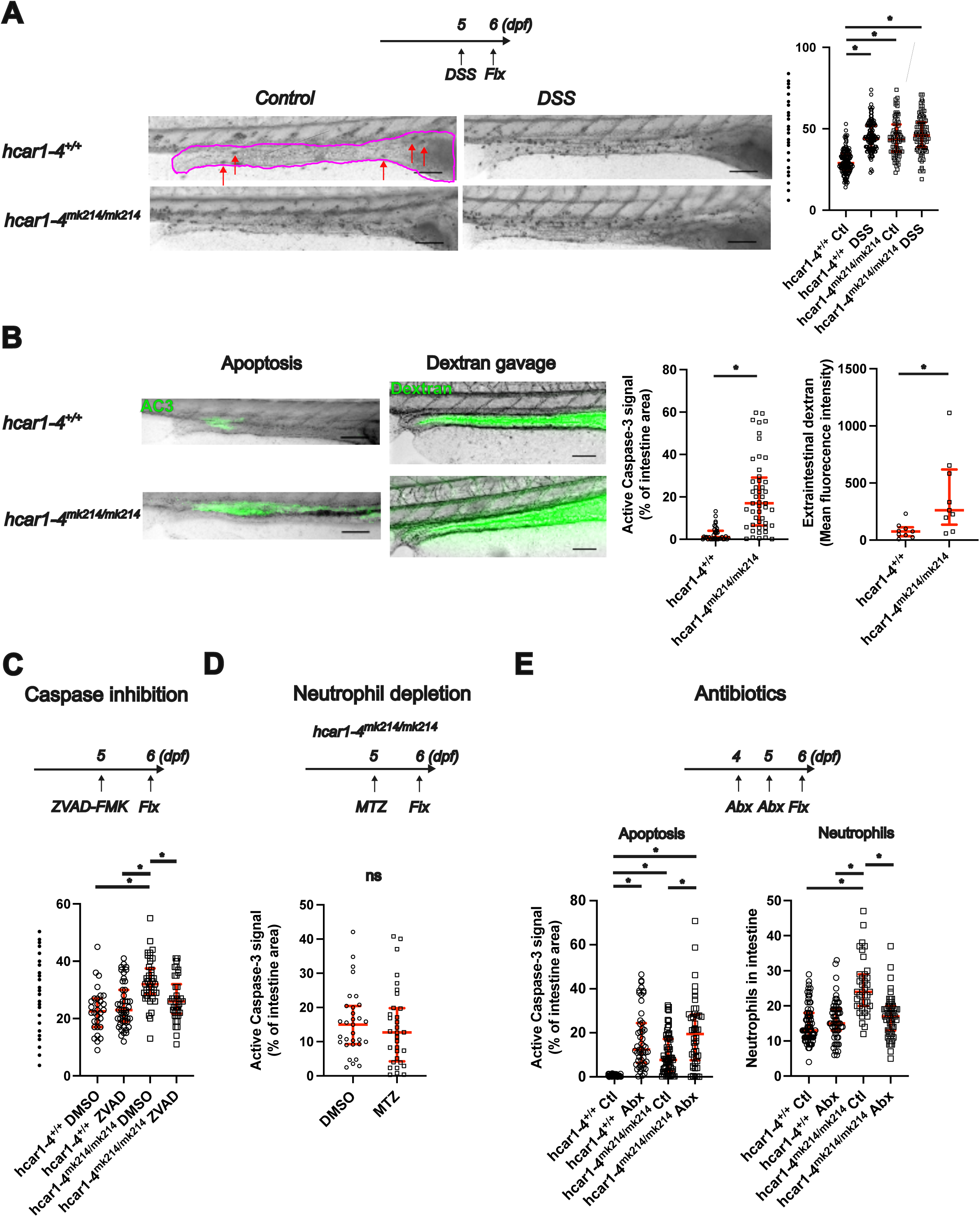
Spontaneous intestinal inflammation in hcar1-4^mk214/mk214^ zebrafish. **(A)** Top: Experimental scheme for dextran sodium sulphate (DSS) colitis model. Left: Representative images of Sudan Black stained larvae (intestine is outlined in magenta, red arrows point to example neutrophils stained by Sudan Black). Note the increased neutrophil counts in control hcar1-4^mk214/mk214^ larvae. Scale bars: 100 µm. Right: Quantification of intestinal neutrophil counts. *: p<0.05, Kruskal-Wallis ANOVA, followed by Dunn’s post hoc test (n=76-124 larvae). **(B)** Left: Representative images of active caspase-3 immunostaining (green) of hcar1-4^+/+^ and hcar1-4^mk214/mk214^ larvae. Scale bars: 100 µm. Middle: Representative images of FITC-dextran (green) gavaged larvae 60 minutes after gavage. Scale bars: 100 µm. Right: Quantification of intestinal active caspase-3 immunostaining. *: p<0.05, Mann Whitney test (n=35 and 48 larvae) and quantification of extraintestinal dextran fluorescence intensity. *: p<0.05, Student’s t-test (n=9-9 larvae). **(C)** Top: Experimental scheme for ZVAD-FMK treatment. Animals were treated for 24 hours with 300 µM ZVAD-FMK or vehicle control. Bottom: Quantification of intestinal neutrophil counts. *: p<0.05, Kruskal-Wallis ANOVA, followed by Dunn’s post hoc test (n=30-46 larvae). **(D)** Top: Experimental scheme for neutrophil ablation. Hcar1-4^mk214/mk214^ larvae expressing the NTR2.0 system in neutrophils [Tg(*lyz*:QF2; 5xQUAS: YFP-NTR2.0)] were treated with 150 µM MTZ or vehicle control for 24 hours. Bottom: Quantification of intestinal apoptosis in control or neutrophil depleted larvae (n=31 and 32 larvae, p=0.43, Mann-Whitney test). **(E)** Top: Experimental scheme for antibiotics treatment. Larvae were treated with ampicillin and kanamycin for 48 hours. Embryo water was switched to fresh water with new antibiotics after 24 hours of treatment. Bottom: Quantification of apoptosis (left) and neutrophil counts (right) in control and antibiotics treated larvae. *: p<0.05, Kruskal-Wallis ANOVA, followed by Dunn’s post hoc test (n=52-61 larvae for the apoptosis assay and 39-69 larvae for the Sudan Black staining)

Given that ROS induces 5-KETE production, 5-KETE may serve as a diffusible oxidative stress ‘danger’ signal that initiates adaptive responses to prevent ROS-induced damage. In this case, *hcar1-4* deficient animals should have an increased sensitivity to ROS-induced damage and cell death. To test this hypothesis, we treated animals with different antioxidants (see scheme in Figure 2A). Scavenging of hydrophilic ROS with Trolox rescued both apoptosis (Figure 2B) and neutrophilic inflammation (Figure 2C) in *hcar1-4* mutant intestines.In contrast, scavenging of hydrophobic ROS with Liproxstatin-1 did not affect apoptosis or neutrophilic inflammation in *hcar1-4* mutant intestines (Supplementary Figure 2A-B). Altogether this suggests that Hcar1-4 suppresses the toxicity of hydrophilic ROS.

**Figure 2.**
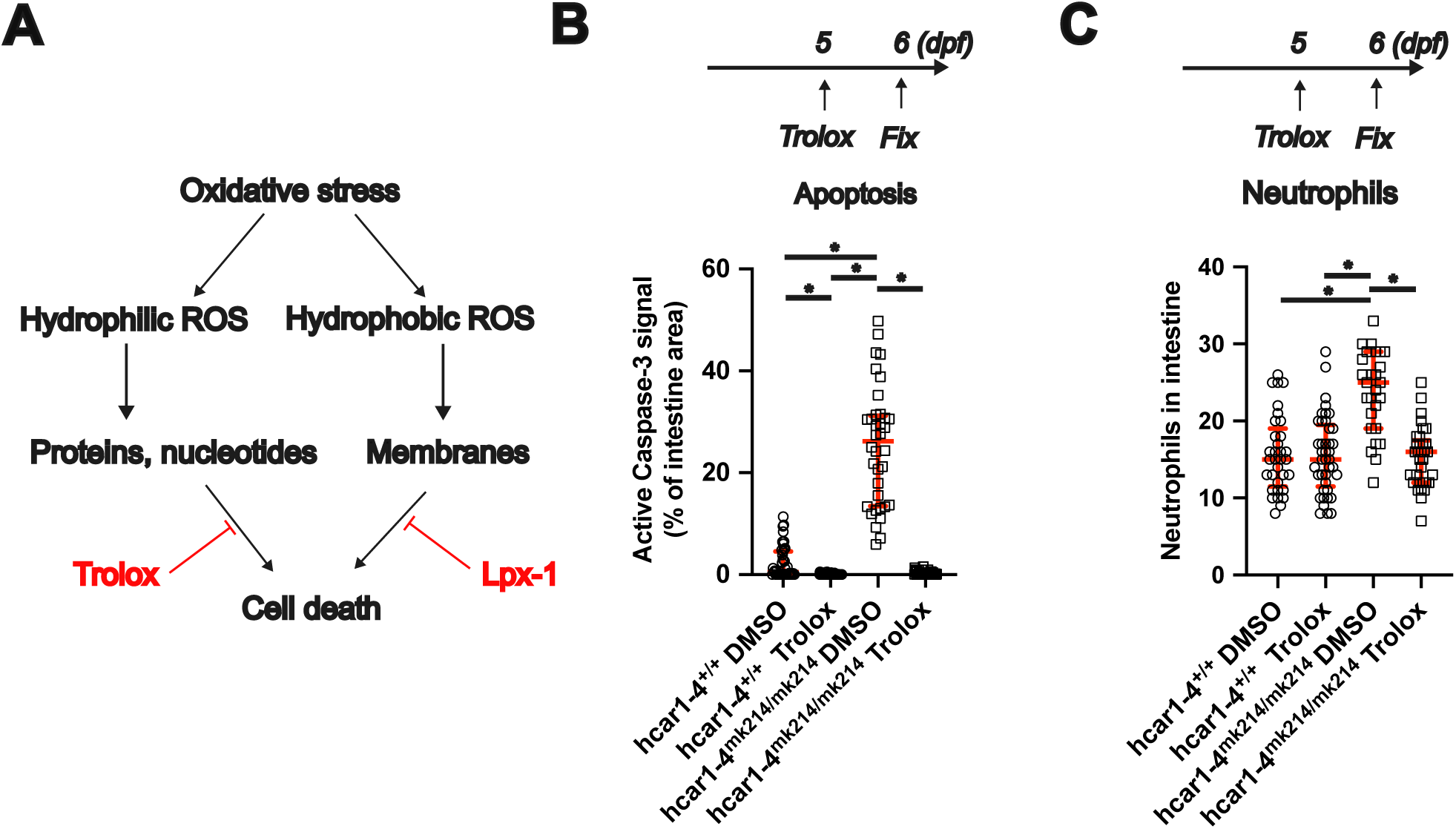
Impaired adaptation to hydrophilic ROS leads to gut epithelial cell apoptosis in hcar1-4^mk214/mk214^ zebrafish. **(A)** Schematic summarizing how oxidative stress induced damage may lead to cell death. Pharmacological tools used to scavenge different free radical types are marked with red. **(B)** Top: Experimental scheme for Trolox treatment. Larvae were treated with Trolox or DMSO control for 24 hours. Bottom: Quantification of apoptosis in control and Trolox treated larvae. *: p<0.05, Kruskal-Wallis ANOVA, followed by Dunn’s post hoc test (n=36-54 larvae). **(C)** Top: Experimental scheme for Trolox treatment. Larvae were treated with Trolox or DMSO control for 24 hours. Bottom: Quantification of neutrophil counts in control and Trolox treated larvae. *: p<0.05, One-way ANOVA, followed by Tukey’s post hoc test (n=27-41 larvae).

To test for phylogenetic conservation of OXER1’s antiapoptotic effects, we silenced the human receptor in monolayer cultures of human Caco-2 cells. OXER1 knockdown (Figure S3A) elevated apoptosis (Figure 3A), which was prevented by Trolox but not liproxstatin-1 treatment (Figure 3B, Supplementary Figure 3B). Pretreatment with physiologically relevant concentrations (1 μM) of 5-KETE blunted H_2_O_2_ induced apoptosis compared to the vehicle control (Figure 3C). Thus, activation of OXER1 increases ROS resilience of intestinal cells across phylae—but how? Our previously published RNA sequencing data [13] revealed decreased NUDT15 expression in *hcar1-4^mk213/mk213^* mutant larvae (Supplementary Figure 3C) while other antioxidant systems remained unchanged by *hcar1-4* knockout. NUDT15 (also known as MTH2) is one of several Nudix hydrolase enzymes (NUDT1, NUDT15 and NUDT18, also known as MTH1-3) involved in the elimination of oxidized purine nucleotides [19]. Silencing of NUDT1, but not NUDT15 increased apoptosis in Caco-2 cells (Supplementary Figure 3D), revealing it as the relevant enzyme in human cells. OXER1 and NUDT1 knockdown both decreased NUDT1 protein expression (Figure 3D, Supplementary Figure 3E) and NUDT1 immunofluorescence staining (Figure 3E, Supplementary Figure 3F). Conversely, pretreatment with 5-KETE enhanced NUDT1 expression (Figure 3F). Notably, the antiapoptotic effect of 5-KETE pretreatment was lost in NUDT1-silenced cells (Figure 3G). Altogether these experiments suggest a phylogenetically conserved mechanism where Nudix hydrolases (Nudt15 and NUDT1) mediate the anti-genotoxic and -apoptotic functions of OXER1 in zebrafish and human cells.

**Figure 3.**
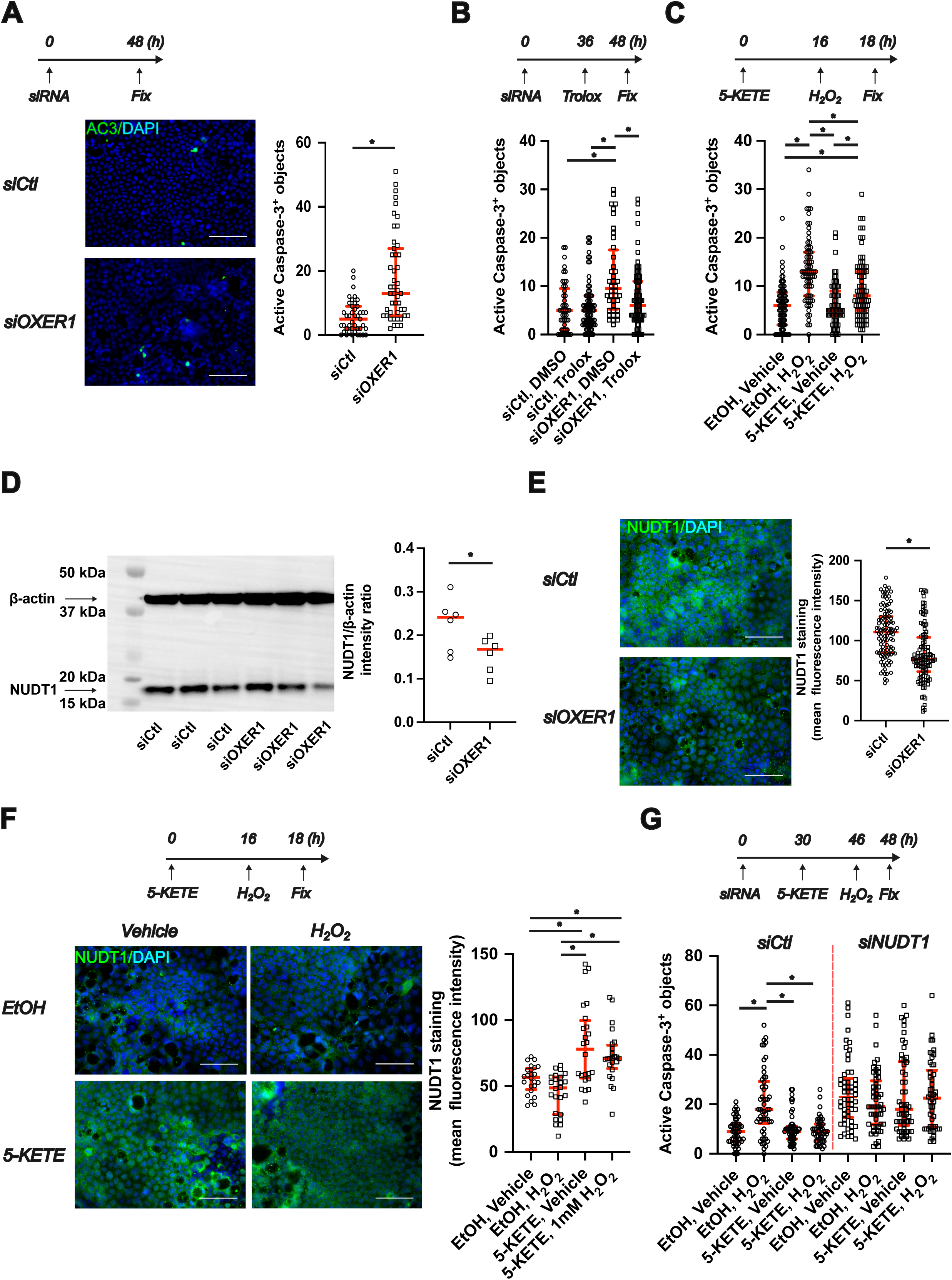
The protective function of OXER1 is conserved in the human intestinal epithelium and mediated by NUDT1. **(A)** Top: Experimental scheme. Caco-2 monolayers were transfected with OXER1 targeting or non-targeting control siRNA. Cells were fixed and used for anti-active caspase-3 immunostaining 48 hours after transfection. Left: Representative microscopic images of anti-active caspase-3 immunostained (green) cells. Nuclei were counterstained with DAPI (blue). Scale bars: 100 µm. Right: Quantification of apoptosis. *: p<0.05, Mann-Whitney test (n=43-47 images from N=2 transfections). **(B)** Top: Experimental scheme. Caco-2 monolayers were transfected with OXER1 targeting or non-targeting control siRNA. Cells were treated with 50 µM Trolox or vehicle control 36 hours after transfection (overnight treatment). Cells were fixed and used for anti-active caspase-3 immunostaining 48 hours after transfection. Bottom: Quantification of apoptosis. *: p<0.05, Kruskal-Wallis ANOVA, followed by Dunn’s post hoc test (n=59-108 images from N=2 transfections). **(C)** Top: Experimental scheme. Caco-2 monolayers were treated with 1 µM 5-KETE or vehicle control overnight. Cells were then challenged with 1 mM H_2_O_2_ or vehicle control for 2 hours. Cells were fixed and used for anti-active caspase-3 immunostaining 48 hours after transfection. Bottom: Quantification of apoptosis. *: p<0.05, Kruskal-Wallis ANOVA, followed by Dunn’s post hoc test (n=69-76 images from N=3 independent experiments). **(D)** Cells were transfected with OXER1 targeting siRNA as in panel A. Whole cell lysates were collected 48 hours after transfection and NUDT1 expression was determined using Western blot. Left: Representative Western blot of OXER1 silenced cells. Blots were probed with anti-NUDT1 and anti-ß-actin antibodies simultaneously, expected molecular weights are marked with arrows. Right: Quantification of NUDT1 band intensity normalized to the ß-actin band intensity *: p<0.05, t-test (n=6-6 samples from N=2 transfections). **(E):** Caco-2 monolayers were transfected with either non-targeting control siRNA or siRNA targeting OXER1. Cells were fixed and used for anti-NUDT1 immunostaining 48 hours after transfection. Left: Representative images of NUDT1 immunostaining (green). Nuclei were stained with DAPI (blue). Right: quantification of NUDT1 immunostaining. *: p<0.05, Mann-Whitney test (n=107 and 108 images from N=2 transfections). **(F)** Top: Experimental scheme. Caco-2 monolayers were treated with 1 µM 5-KETE or vehicle control overnight. Cells were then challenged with 1 mM H_2_O_2_ or vehicle control for 2 hours. Cells were fixed and used for anti-NUDT1 immunostaining 48 hours after transfection. Left: Representative images of anti-NUDT1 immunostaining. Right: Quantification of NUDT1 staining. *: p<0.05, Kruskal-Wallis ANOVA, followed by Dunn’s post hoc test (n=24-24 images from N=1 independent experiment). (**G)** Top: Experimental scheme. Caco-2 monolayers were transfected with NUDT1 targeting or non-targeting control siRNA. 36 hours after transfection, cells were treated with 1 µM 5-KETE or vehicle control overnight. Cells were then challenged with 1 mM H_2_O_2_ or vehicle control for 2 hours. Cells were fixed and used for anti-active caspase-3 immunostaining 48 hours after transfection. Bottom: Quantification of apoptosis: *: p<0.05, Kruskal-Wallis ANOVA, followed by Dunn’s post hoc test (n=49-54 images from N=2 transfections). All NUDT1 silenced groups were significantly different from the siCtl EtOH,Vehicle; 5-KETE, Vehicle; 5-KETE, H_2_O_2_ groups. The siCtl,EtOH,H_2_O_2_ group was not significantly different from any of the NUDT1 silenced groups.

To further test this idea *in vivo,* we used two complementary approaches: First, we knocked out Nudix hydrolases in WT zebrafish with an F0 CRISPR approach (See Figure 4A for experimental scheme and Supplementary Figure 4 for details), and second, we overexpressed Nudix hydrolases in the gut epithelium of *hcar1-4^mk214/mk214^* zebrafish using the intestinal epithelium specific *cldn15la* promoter (see Figure 4B for validation of the promoter and experimental scheme). F0 CRISPR *of nudt15* increased apoptosis in the intestine (Figure 4C), while F0 CRISPR of *nudt1* had no effect (Supplementary Figure 5A). F0 CRISPR larvae for both *nudt1* and *nudt15* were grown to sexual maturity without obvious defects. F0 CRISPR of *nudt15* increased intestinal neutrophil counts in hcar1-4^+/+^ larvae but had no effect in *hcar1-4^mk214/mk214^* larvae (Figure 4D). As the *hcar1-4* and *nudt15* deficiency phenotypes were not additive, both proteins are likely part of the same genetic pathway. Expression of zebrafish NUDT15 or human NUDT1 in the gut epithelium using the intestinal epithelium specific *cldn15la* promoter in *hcar1-4^mk214/mk214^* animals reduced neutrophil counts in the intestine (Figure 4E).

**Figure 4.**
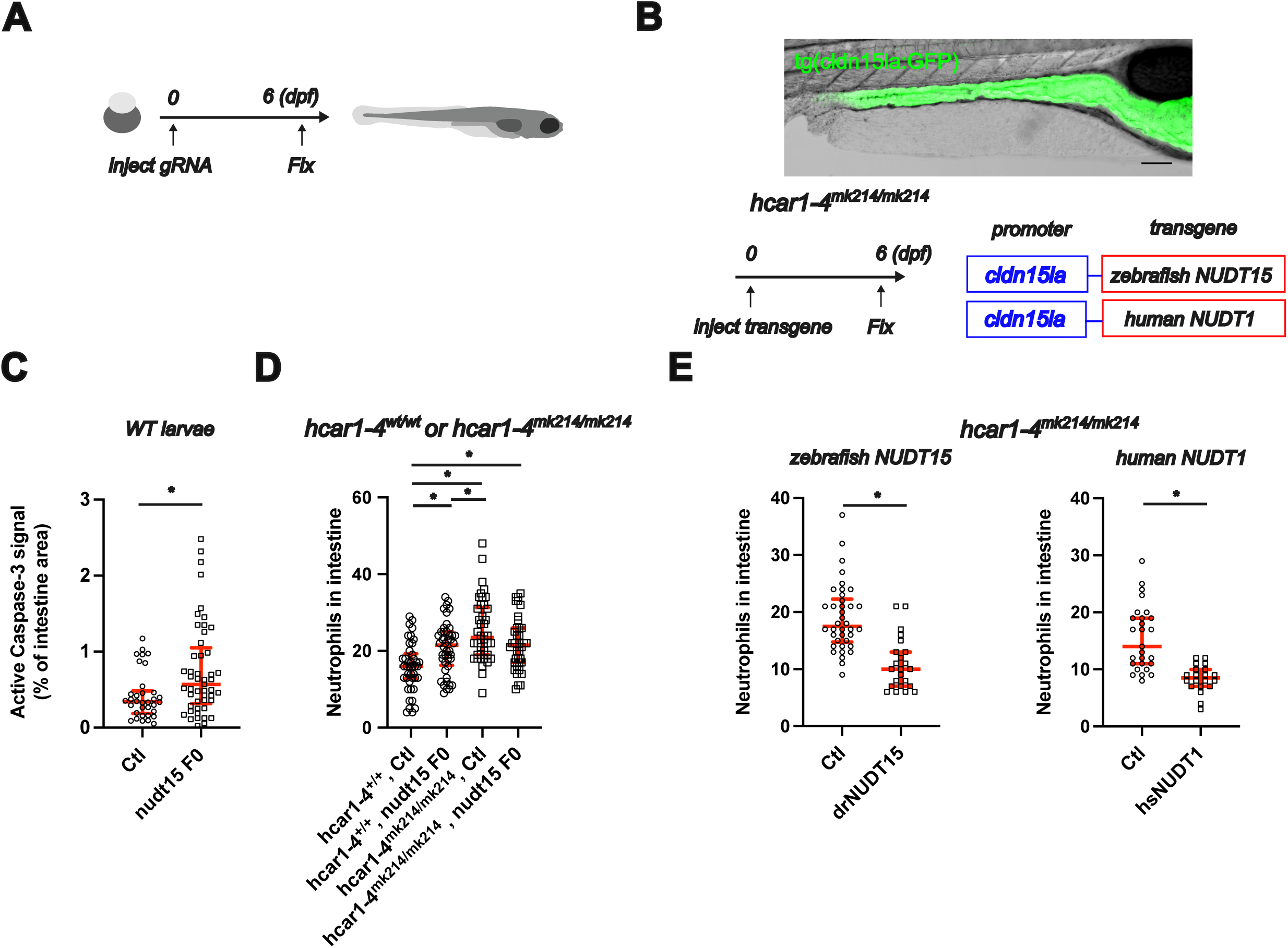
The hcar1-4^mk214/mk214^ phenotype *in vivo* is mediated by NUDT15 and rescued by NUDT overexpression. **(A)** Experimental scheme for the F0 CRISPR experiments. Zebrafish embryos (wild-type in Figure 4C, hcar1-4^+/+^ and hcar1-4^mk214/mk214^ in Figure 4D) were injected with Cas9 and guide RNAs for *nudt15* at the one cell stage. Larvae were fixed at 6dpf and used for anti-active caspase-3 immunostaining or Sudan Black staining. Larval zebrafish graphic was sourced from Scidraw.io (courtesy of Luigi Petrocco, doi.org/10.5281/zenodo.3926123) **(B)** Top: representative image showing expression of GFP in the intestinal epithelium of Tg(*cldn15la*:QF2 xQUAS:GFP) fish. Bottom: Experimental scheme. One-cell stage hcar1-4^mk214/mk214^ zebrafish embryos were injected with Tol2 transposase mRNA and transgenesis plasmids for intestinal epithelial expression of zebrafish NUDT15 or human NUDT1 (driven by the *cldn15la* promoter). F0 transgenic animals were identified based on selection marker expression. Larvae were fixed at 6dpf and used for Sudan Black staining. **(C)** Quantification of apoptosis in control or *nudt15* F0 CRISPR larvae *: p<0.05, Mann-Whitney test (n= 34 and 49 larvae). **(D)** Quantification of intestinal neutrophil counts in control or *nudt15* F0 CRISPR larvae for both hcar1-4^+/+^ and hcar1-4^mk214/mk214^ larvae. *: p<0.05, One-way ANOVA, followed by Tukey’s *post hoc* test (n=38-44 larvae). **(E):** Quantification of intestinal neutrophil counts for hcar1-4^mk214/mk214^ larvae expressing zebrafish NUDT15 (left) or human NUDT1 in the intestinal epithelium. *: p<0.05, Mann-Whitney test (n=23-38 larvae for zebrafish NUDT15) or Student’s t-test (n=20-27 larvae for human NUDT1).

In conclusion, our data reveal a conserved, non-immune function of 5-KETE signaling. Besides its previously described chemotactic functions, our data show that the OXER1 pathway acts anti-apoptotic on the intestinal epithelium through enhancing the redox protection of the cellular nucleotide pool *in vivo* [19, 20]. Thus, OXER1 signaling orchestrates microbicidial leukocyte responses via increasing the ROS resilience of the affected epithelium. By limiting damage of the “bystanding” tissue, hosts can increase the “effective selectivity” of the otherwise unselectively toxic, innate defense response. Our data highlight ROS mucosal adaptation as an integral part of innate immune signaling, and suggests that not an “arms race” but a “shields race” determines efficiacy of at least some innate defense mechanisms.

Beyond its conceptual novelty, our work has potential therapeutic implications for mucosal inflammation and tumorigenesis. 5-KETE levels are elevated in the unaffected mucosa of patients with inflammatory bowel disease compared to inflamed mucosa from the same patients and samples from healthy controls, aligning with the idea that oxoeicosanoid signaling may protect the human intestine [21]. Our work suggests that OXER1 signaling may have antimutagenic effects by protecting host cells from ROS damage. Interestingly, a frequent frameshift mutation (S78Pfs*64 or S78Vfs*61) leading to a truncated, non-functional OXER1 has been reported in human gastrointesinal cancers [22]. The discovery of the redox-sensitive, 5-KETE-producing enzyme 5-HEDH in our companion paper (Ma et al., 2025) now opens the possibility to probe some physiological aspects of this vastly understudied redox signaling pathway also in rodents. Our current approach highlights the benefits of combining alternative model systems and human cell lines to illuminate “blind spots” of mouse genetics, while ensuring relevance to human biology. A better understanding of 5-KETE signaling should open up new avenues for mucoprotective therapies that protect barrier tissues against exessive oxidative stress.

## Author contributions

ML and PN conceived the project. ML performed most of the experiments alone or with assistance from SP (initial *in vivo* colitis experiments, creating transgenic lines) and MQ (initial *in vitro* experiments). YM generated fish lines and helped perform the F0 CRISPR experiments. ZG generated the tg(*lyz*:NTR2.0) fish line and established protocols for chemogenetic ablation.

## Acknowledgements

The research has been funded by the NIH grant R35GM140883 and a BRIA award to PN, as well as a Marie-Josée Kravis Women in Science Endeavor Postdoctoral Fellowship to YM, an Experimental Immunooncology Scholars fellowship and a Crohn’s Colitis Foundation Research Fellows Award (Award #:1268111) to ML. Core facility services were in part funded by the NIH/NCI Cancer Center Support grant P30CA008748. We would like to thank Péter Enyedi, Balázs Enyedi, Tobias C. Walther and Tim Mitchison for valuable comments on the manuscript.

## Methods

### General zebrafish procedures

Adult *casper* zebrafish were maintained as described as previously [23] and subjected to experiments according to institutional animal healthcare guidelines with the approval of the Institutional Animal Care and Use Committee (IACUC). Generation and genotyping of *alox5a* and *hcar1-4* mutants used in this study were described previously [13, 17]. Both the neutrophil and apoptosis phenotypes were more robust in the *hcar1-4^mk214/mk214^* line (this line has a large truncation and a mutation at an upstream part of the *hcar1-4* coding region, whereas the *mk213* line is mutated at a more downstream genomic region), so we used this line for the majority of the experiments. Zebrafish larvae were maintained in standard hypotonic E3 medium (5 mM NaCl, 0.17 mM KCl, 0.33 mM CaCl_2_, 0.33 mM MgSO_4_) until used for experiments at 6 dpf. At this developmental stage, sex cannot be determined and is unlikely to influence the biological processes under study.

### Larval zebrafish treatments

For the chemical colitis experiments, dextran sodium sulfate was obtained from Sigma (42867). On the day of the experiment, a 10 w/v% stock of DSS was made in embryonic E3 media as described previously [16]. DSS was further diluted to a working concentration of 0.25% w/v in E3. 5dpf larvae (WT or mutant) were randomly distributed to E3 with or without DSS for 24 hours. The animals were euthanized and fixed using 4% formaldehyde in PBS or PBSTX before further processing. ZVAD-FMK was obtained from ApexBio (A1902) dissolved in DMSO and stored as a 300 mM stock solution at −20°C. The drug was used at a working concentration of 300 µM in E3 as described by others previously [24, 25]. 5dpf larvae were randomly distributed to E3 with ZVAD-FMK or 0.1% DMSO for 24 hours. The animals were euthanized and fixed using 4% formaldehyde in PBS before further processing. Metronidazol (MTZ) was obtained from Sigma (PHR1052-1G) dissolved in DMSO and stored as a 100 mM stock solution at −20°C. The drug was used at a working concentration of 150 µM in E3 as described by us and others [18, 26]. 5dpf *hcar1-4*^mk214/mk214^ tg(*lyz*:QF2 x 5xQUAS:NTR2.0) larvae were randomly distributed to E3 with MTZ or 0.15% DMSO for 24 hours. The animals were euthanized and fixed using 4% formaldehyde in PBS with 0.05% Triton-X (PBSTX) before further processing. Ampicillin (BP1760-25) and Kanamycin (BP906-5) were obtained from Fisher dissolved in distilled water and stored at −20°C. Antibiotics were used at a working concentration of 100 µg/ml and 30 µg/ml in E3 for ampicillin and kanamycin, respectively. 4 dpf larvae were randomly distributed to E3 with either Ampicillin/Kanamycin or 0.2% distilled water. Larvae were transferred to fresh E3 with or without antibiotics after 24h. Animals were euthanized and fixed using 4% formaldehyde in PBS or PBSTX before further processing. Trolox was obtained from Selleck chemicals (S3665), dissolved in DMSO and stored as a 50 mM stock solution at −20°C. The drug was used at a working concentration of 50 µM as described previously by others [27]. Liproxstatin-1 (SML1414-5MG) was obtained from Sigma Aldrich dissolved in DMSO and stored as a 100 mM stock solution at −20°C, and used at a working concentration of 20 µM as described previously [17].

### Whole mount zebrafish staining

#### Sudan Black

Intestinal neutrophils were visualized by Sudan Black staining as described by us and others previously [13, 17, 28]. In brief, Larvae were fixed in 4% formaldehyde in PBS overnight at 4 °C, washed 3 times with PBS and stained with Sudan Black (Sigma Aldrich, 199664) for 1 hour. Larvae were washed 3 times with 70% ethanol and rehydrated in PBS with 0.1% Tween (PBST) for 5 minutes. Larvae were then treated with depigmentation solution (1% KOH, 1% H_2_O_2_ in water) for 5 minutes followed by 2 PBST washes before imaging.

#### Active caspase-3 immunostaining

Active caspase-3 was visualized using the protocol described by others with minor modifications [29]. Larvae were fixed in 4% formaldehyde in PBSTX overnight at 4°C. Larvae were then washed with PBSTX and blocked for 2 hours at room temperature in a blocking solution (PBSTX supplemented with 2 mg/ml bovine serum albumin (BSA) and 1% DMSO). Primary antibody (anti-active caspase-3 (BD Biosciences Product: 559565)) was diluted 1:100 in blocking solution and larvae were incubated overnight at 4°C. Larvae were washed with PBSTX and incubated with secondary antibody conjugated with Alexa Fluor 488 or 555 (Abcam, ab150077 or ab150078, respectively) diluted 1:1.000 in blocking solution overnight at 4℃. Finally, larvae were washed with PBSTX and mounted for imaging.

### Microgavage of larval zebrafish

Microgavage was performed as described previously by others [30]. 4 kDa molecular weight FITC labeled dextran (Sigma-Aldrich FD4-100MG) was dissolved in PBS to obtain a 10 mg/ml solution. This solution was then loaded into a microinjector needle. Before gavage, 6 dpf zebrafish larvae were anesthetized with 0.2 mg/ml MS-222 (Fisher Scientific, NC0589195) dissolved in E3. After verifying proper depth of anesthesia, the larvae were positioned laterally, ventral side toward the vertical wall of an agarose mold. Afterward, the needle was gently maneuvered down the esophagus and into the anterior intestine. The material of interest was then injected into the lumen (typically, volumes of 5-10 nl were used). The needle was then gently but firmly removed from the animal. Larvae were then transferred to E3 to recover from the anesthesia and kept in E3 for 60 minutes and then used for imaging experiments.

### F0 CRISPR gene perturbation approach for *nudt1* and *nudt15*

F0 CRISPR targeting of *nudt1* and *nudt15* was performed as described previously [13]. Potential gRNA target sites were identified using the web program CHOPCHOP (http://chopchop.cbu.no/index.php). All gRNA target sequences are listed in Supplementary Table S1. Two independent gRNAs for *nudt1* (ENSDARG00000030573) and two independent gRNAs for *nudt15* (ENSDARG00000113961) were designed and ordered from IDT (for sequences see Supplementary Table S1). These target-specific Alt-R crRNA (Crispr RNA) and common Alt-R tracrRNA (Trans-Activating Crispr RNA, IDT 1072532) were dissolved in nuclease-free duplex buffer (IDT; 11-01-03-01) to obtain a 100 mM stock solution. To prepare the crRNA:tracrRNA duplex, equal volumes of 100 mM Alt-R crRNAs (1:1) and 100 mM Alt-R tracrRNA stock solutions were mixed to obtain a final concentration of 10 mM each. The mixture was heated to 95°C for 5 min and then cooled down to room temperature. ALT-R S.p. Cas9 Nuclease (IDT, 1081058) was diluted to 1 mg/mL with Cas9 working buffer (20 mM HEPES, 150 mM KCl, pH 7.5). The Cas9-gRNAs ribonucleoprotein complex was assembled by combining the crRNA:tracrRNA duplex and Cas9 protein solutions at a 1:1 ratio, incubated at 37℃ for 5 min and then placed at room temperature. 2.3 nL of the Cas9-gRNAs ribonucleoprotein complex solution was injected into the cytoplasm of one-cell-stage zebrafish embryos. At 2.5-3 dpf, genomic DNA was isolated from individual F0 larvae for genotyping. Each specific target sequence of interest was PCR amplified from the genomic DNA samples with the primers listed in Table S1. Next, PCR products were incubated with the FastDigest (Thermo) restriction enzymes MvaI (FD0554) for *nudt15*-g1 Cfr10I (FD0184) for *nudt15*-g2, BstXI (FD1024) for *nudt1*-g1, or in the case of *nudt1*-g2, BtsI-v2 (NEB, R0667S) for at least 2 h at 37°C. Then, samples were separated by agarose gel electrophoresis (see Table S1 for primers) and guide efficiency was evaluated (Supplementary Figure 4).

### Plasmid construction

Total RNA was isolated from Caco-2 cells or 35-40 6dpf *hcar1-4*^+/+^ larvae using Trizol (ThermoFisher 15596026) and 1 µg was used as a template for cDNA synthesis using the Revertaid First Strand cDNA synthesis kit (Thermo Scientific K1622). The coding sequence of human NUDT1 and zebrafish NUDT15 were subcloned into a pME-MCS backbone between the Bsu15I and XbaI sites using T4 DNA ligase (NEB M0202S). The sequence fidelity of the resulting pME-hsNUDT1 and pME-drNUDT15 plasmids was confirmed with Sanger sequencing. The *cldn15la*:QF2, *cldn15la*:hsNUDT1 and *cldn15la*:drNUDT15 plasmids were created with Gateway cloning (LR Clonase II enzyme, Thermofisher 11791020) using the Tol2kit system as previously described [31], recombining the *cldn15la* promoter, the QF2 transcriptional activator,hsNUDT1 or drNUDT15 and a SV40 polyadenylation site into the pDestTol2CG2 (for *cldn15la*:QF2) or the pDestTol2CR destination vector (for *cldn15la*:drNUDT15 or *cldn15la*:hsNUDT1). The pDestTol2CR vector was a gift from Balázs Enyedi [32] and the p5E-0.349cldn15la plasmid was a gift from John Rawls (Addgene plasmid # 125026 ; http://n2t.net/addgene:125026; RRID:Addgene_125026) [33].

### Transgenesis

A solution containing 25 pg of the *cldn15la*:QF2 and 25 pg Tol2 transposase mRNA into the cytosol of one-cell stage wild-type embryos. For the rescue experiments in *hcar1-4^mk214/mk214^* animals, a similar mixture of *cldn15la*:hsNUDT1 or *cldn15la*:drNUDT15 plasmid and Tol2 transposase mRNA was injected into the cytosol of one-cell stage *hcar1-4^mk214/mk214^* embryos. Injected larvae with cardiac green fluorescent protein (GFP, for *cldn15la*:QF2) or red fluorescent protein (mKate2, for *cldn15la*:hsNUDT1 or *cldn15la*:drNUDT15) expression were selected 24 hours post-fertilization. Transgenic embryos with Nudix hydrolase expression were used for experiments in the F0 generation, while *cldn15la*:QF2 embryos were grown to adulthood and used to establish a stable transgenic line. The GFP expressing transgenic larvae shown in Figure 4B were obtained by crossing *cldn15la*:QF2 and QUAS:eGFP[26] animals and selecting fish showing GFP expression in the intestine.

### Cell culture

Caco-2 cells were obtained from ATCC. Cells were maintained in EMEM (ATCC, 30-2003) supplemented with Pen-Strep (Corning, MT30002CI) and 20% heat-inactivated FBS (Sigma, F2442). For experiments, cells were trypsinized and seeded into 8-well precoated imaging dishes (ibidi, 80826) at a density of 8000 cells/well and grown to confluence for experiments.

### Transfection of siRNA

A pool of 3-4 independent siRNAs targeting OXER1 (L-005741-00-0005), NUDT1 (L-005218-00-0005), NUDT15 (L-020621-01-0005) or non-targeting controls (D-001810-10-05) were ordered from Horizon Discovery, reconstituted in 1x siRNA buffer ( 5x siRNA buffer B-002000-UB-100, Horizon Discovery diluted to 1x with RNAse free water B-003000-WB-100, Horizon Discovery) at a concentration of 25 µM and stored at −20 °C. Cells were transfected with siRNA using Dharmafect 1 transfection reagent (T-2001-02) according to the manufacturer’s instructions. In brief, 5 µl of siRNA and 2.5 µl of Dharmafect 1 transfection reagent were diluted separately in 100 µl complete EMEM each, incubated at room temperature for 10 minutes and then mixed and incubated at room temperature for 20 minutes. Afterwards, 800 µl of complete EMEM was added and the mixture was added to the cells (250 µl/ well of an ibidi 8-well dish). Cells were assayed 48 hours after transfection.

### Drug treatments in cell culture

Trolox was used at a working concentration of 50 µM and Liproxstatin-1 was used at a working concentration of 20 µM in serum-free EMEM. For experiments with 5-KETE, the lipid was obtained as an ethanol-based solution (Cayman Chemical, 34250) and was used at a working concentration of 1 µM in serum-free EMEM. Cells were treated with the compounds or the appropriate vehicle control overnight. For H_2_O_2_ treatment experiments, a 1 M stock solution was diluted from 30% H_2_O_2_ on the day of the experiment. Cells were treated with 1 mM H_2_O_2_ or vehicle control diluted in serum-free EMEM for 2 hours.

### Western blotting

Caco-2 cells were grown in 6-well plates and transfected with siRNA using the protocol detailed above using 1 ml transfection mixture per well. Cells were washed once with ice-cold PBS and harvested 48 hours after transfection in RIPA buffer (Cell Signaling, #9806) containing protease inhibitors (Roche, 11836153001). Lysates were centrifuged at 18,000 *g* for 10 minutes at 4°C and protein content was determined using the BCA assay (Thermo Scientific, A65453) according to manufacturer’s instructions. Lysates containing 10 µg protein were treated with reducing agent (50 mM DTT, Thermo, NP0009) and sample loading buffer (NP0007, Thermo), boiled for 10 minutes and subjected to SDS-PAGE on precast gradient gels (NP0322, Thermo) at 120V for 120 minutes. Proteins were transferred to nitrocellulose membranes using standard procedures. Membranes were blocked in 5% skim milk dissolved in TBS containing 0.1% Tween-20 (TBS-T) for 2 hours at RT. Primary antibodies were diluted in TBS-T containing 3% BSA and rocked at 4 ℃ overnight (anti-NUDT1 (abcam, ab200832), 1:5000; anti-ß-actin (Sigma, A5441), 1:5000). Afterwards, blots were washed 5 times with TBS-T at RT and incubated with secondary antibodies conjugated to HRP (anti-mouse (Cell Signaling, 7076S), 1:4000, anti-rabbit (Cell Signaling, 7074S), 1:2000) diluted in TBS-T containing 2.5% skim milk for 1-2 hours at RT. Blots were washed 5 times in TBS-T and signals were detected using an enhanced chemiluminescent detection reagent according to manufacturer’s instructions (Cytiva, RPN2235).

### RT-PCR of Caco-2 cells

Total RNA was isolated from control or OXER1 silenced Caco-2 cells using Trizol and 1 µg was used as a template for cDNA synthesis using the Revertaid First Strand cDNA synthesis kit. PCR was carried out using Phusion polymerase with previously published primers targeting OXER1 and ß-actin (for sequences see Table S1)[34, 35].

### Immunofluorescent staining of Caco-2 cells

Immunofluorescent staining of fixed Caco-2 cells was performed as described previously with some modifications [36]. In brief, cells were fixed with 4% formaldehyde in PBS at RT for 15 minutes, washed 3 times with PBS, permeabilized with PBSTX for 15 minutes and blocked with 5% goat serum (Jackson Immunoresearch, 005-000-121) in PBS at RT for 1 hour. Cells were incubated with primary antibodies (anti-active caspase-3 1:200, anti-NUDT1, (Abcam, ab200832) 1:100) diluted in blocking solution at RT for 1.5 hours. Cells were washed 3 times with PBS and then incubated with secondary antibodies (1:500) diluted in blocking solution at RT for 30 minutes. Cells were washed twice with PBS and stained with DAPI (1:5000) in PBS for 15 minutes. Cells were 2 more times with PBS before imaging.

### Microscopy

#### Imaging Sudan Black staining, gavaged dextran and active caspase-3 immunostaining in larvae

Larvae were embedded in 200 µl 1% low-melting agarose in a glass-bottom dish (MatTek Corporation; P35G-1.5-20C). For the experiments with live animals, E3 media with tricaine was added to the top of the agarose after it solidified to prevent desiccation of the sample and maintain anesthesia over the course of the imaging. Transmitted light images were acquired at room temperature (∼25 °C) using NIS-Elements (Nikon) on a Nikon Eclipse Ti microscope equipped with a 10×, plan-apochromat NA 0.45 air objective lens, and an Andor Clara charge-coupled device (CCD) camera. Alexa 488 and FITC fluorescence was excited using an LED light source (Lumencor 7; Light Engine) with an internal 475/28 bandpass filter, and Alexa 555 fluorescence was excited using an internal 549/15 bandpass filter. The emission of Alexa 488 and FITC was sampled using a 525/50 bandpass filter (Chroma; 525/50m), while the emission of Alexa555 was detected using a 632/60 bandpass filter (Chroma; ET632/60m). Z-stacks were acquired with a 5 µm step size with a total imaging depth of 80 µm.

#### Confocal imaging of cldn15la:GFP larvae

Larvae were anesthetized and embedded in 200 µl 1% low-melting agarose in a glass-bottom dish (MatTek Corporation; P35G-1.5-20C). Images were acquired at room temperature using NIS-Elements on an inverted Nikon Eclipse Ti microscope equipped with a 20x air objective, a motorized stage, a Yokogawa CSU-W1 Spinning Disk unit, a Photometrics Prime BSI Scientific CMOS camera (2x2 binning). Fluorophores were excited using a 488 nm diode laser (LUN-F 100-240V∼50/60Hz, Nikon Instruments Inc). Emitted light was collected using a 525/36 bandpass filter (Chroma Technology Corp., 89100bs). The imaging plane 1331 μm x 1331 μm x 100 μm) was collected at 5 μm z step size to result in 21 slices.

#### Imaging fixed Caco-2 cells

Caco-2 cells were imaged on the same setup detailed previously for imaging active caspase-3 immunostained zebrafish larvae. DAPI fluorescence was excited using an LED light source (Lumencor 7; Light Engine) with an internal 390/18 bandpass filter and detected using a 460/50 bandpass filter (Chroma; ET460/50m). Images were acquired in a single z-plane.

### Image analysis

#### Zebrafish experiments

##### Sudan Black

Image stacks were imported into ImageJ and used to generate minimum intensity projections. Neutrophils were counted manually in the intestine area.

##### Dextran gavage

Image stacks were imported into ImageJ and used to generate maximum intensity projections. The projections were background subtracted and rectangular regions of interests were applied ∼30-50 µm from the intestine with the longer side of the rectangle parallel to the axis of the intestine. Results are given as mean fluorescence intensity in the ROI as quantified in Fiji.

##### Active caspase-3 immunostaining

Transmitted light image stacks were imported into Fiji and used to generate minimum intensity projections, which were used to label the intestine as a ROI for each animal. Image stacks of the active caspase-3 signal (Alexa 488 or Alexa555) were imported into ImageJ and used to generate maximum intensity projections. The projected images were then background subtracted. Images were thresholded using the MinError method in Image J, converted to binary images and segmented using the watershed algorithm in Fiji. Particles smaller than 5 µm^2^ were omitted from the analysis. Results were given as % positive area of the total intestinal area.

#### Caco-2 cell experiments

##### NUDT1

The abundance of the fluorescent signal was determined as the mean fluorescence intensity of the whole background subtracted image. Background was determined using images of cells which were not stained with the primary antibody.

##### Active caspase-3

Images were background subtracted, thresholded using the MinError method, converted to binary images and segmented using the watershed algorithm in Fiji. Particles smaller than 5 µm2 were omitted from the analysis. Results are given as a number of positive particles (similar results were obtained using % of total area).

#### Statistical analysis

Statistical analysis was performed using GraphPad Prism. Outliers were identified and removed using the Rout method in Prism. The normality of the cleaned data was checked using the Shapiro-Wilk test. Data with normal distributions were compared with the Student’s t-test (in the case of equal variances, as determined by the F-test) or Welch’s t-test (in the case of unequal variances, as determined by the F-test) (2 groups) or One-Way ANOVA followed by Tukey’s *post hoc* test (at least 3 groups), while data that did not have a normal distribution were compared with the Mann-Whitney test (2 groups) or the Kruskall-Wallis ANOVA followed by Dunn’s *post hoc* test (3 or more groups). Differences were considered statistically significant at p<0.05. Statistical analysis for all data in this study is summarized in Supplementary Table S2.

## Figure Legends

**Figure S1.**
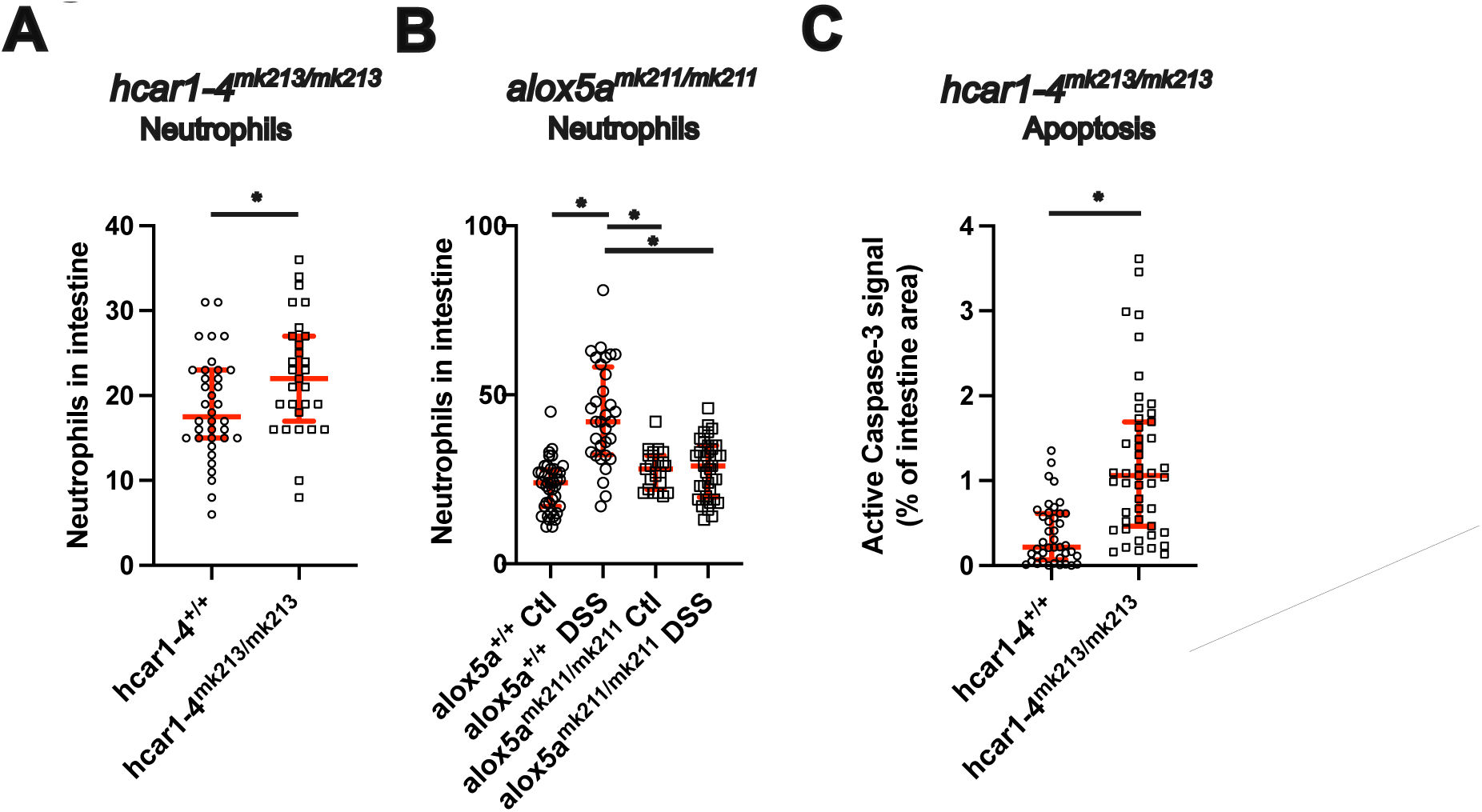
Extended data supporting Figure 1. **(A)**: Quantification of intestinal neutrophil counts in 6dpf hcar1-4^+/+^ or hcar1-4^mk213/mk213^ larvae, *: p<0.05, Student’s t-test (n=29 and 36 larvae). **(B)**: Quantification of intestinal neutrophil counts in 6dpf alox5a^+/+^ or alox5a^mk211/mk211^ larvae after overnight DSS treatment. For the experimental scheme see Figure 1A. *: p<0.05, One-way ANOVA, followed by Tukey’s post hoc test (n=23-40 larvae). **(C)**: Quantification of intestinal anti-active caspase-3 immunostaining in 6dpf hcar1-4^+/+^ or hcar1-4^mk213/mk213^ larvae. *: p<0.05, Mann Whitney test (n=41 and 47 larvae)

**Figure S2.**
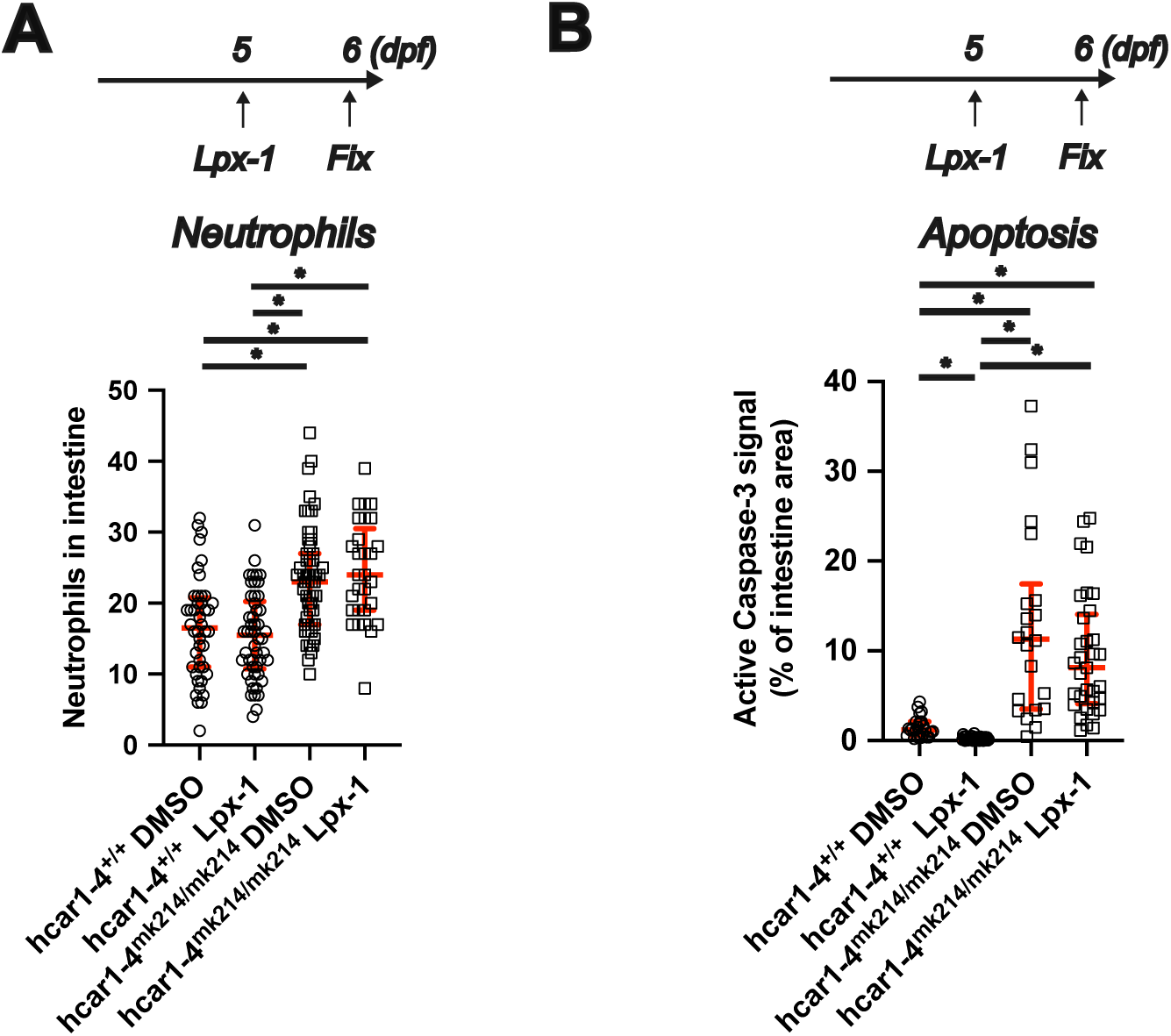
Extended data supporting Figure 2. **(A)**: Top: Experimental scheme for Liproxstatin-1 treatment. Larvae were treated with 20 μM Liproxstatin-1 (Lpx-1) or DMSO control for 24 hours and fixed at 6dpf for further processing. Bottom: Quantification of neutrophil counts in control and Lpx-1 treated larvae. *: p<0.05, One-way ANOVA, followed by Tukey’s post hoc test (n=29-55 larvae).**(B)** Top: Experimental scheme for Liproxstatin-1 treatment. Larvae were treated with Liproxstatin-1 (Lpx-1) or DMSO control for 24 hours and fixed at 6dpf for further processing. Bottom: Quantification of intestinal anti-active caspase-3 immunostaining in control and Lpx-1 treated larvae. *: p<0.05, Kruskal-Wallis ANOVA, followed by Dunn’s post hoc test (n=22-43 larvae).

**Figure S3.**
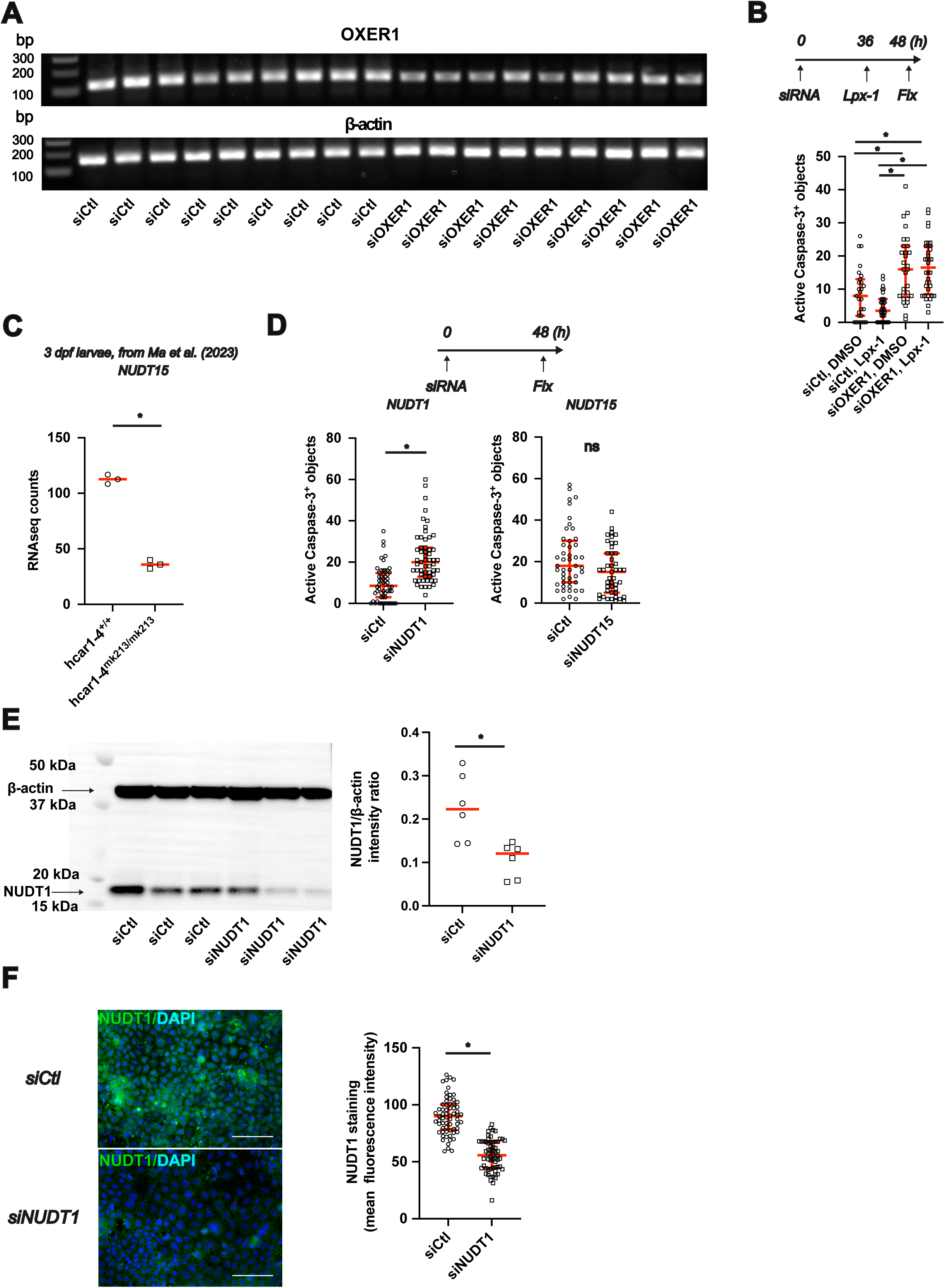
Extended data supporting Figure 3. **(A)** Caco-2 monolayers were transfected with OXER1 targeting or non-targeting control siRNA. Total RNA was isolated 48 hours after transfection and used for cDNA synthesis. Semiquantitative PCR was performed using primers specific for OXER1 (top) and ß-actin (bottom). **(B)** Top: Experimental scheme. Caco-2 monolayers were transfected with OXER1 targeting or non-targeting control siRNA. Cells were treated with 20 µM Lpx-1 or vehicle control 36 hours after transfection (overnight treatment). Cells were fixed and used for anti-active caspase-3 immunostaining 48 hours after transfection. Bottom: Quantification of apoptosis. *: p<0.05, Kruskal-Wallis ANOVA, followed by Dunn’s post hoc test (n=27-36 images from N=1 transfection). **(C)**: mRNASeq counts for NUDT15 from 3dpf hcar1-4^+/+^ and hcar1-4^mk213/mk213^ larvae. Replotted from [13], source data deposited in GEO under number GSE201604. **(D)**: Top: Experimental scheme. Caco-2 monolayers were transfected with either non-targeting control siRNA or siRNA targeting NUDT1 or NUDT15. Cells were fixed and used for anti-active caspase-3 immunostaining 48 hours after transfection. Left: Quantification of apoptosis after NUDT1 silencing. *: p<0.05, Mann-Whitney test (n=55-56 images from N=2 transfections). Right: Quantification of apoptosis after NUDT15 silencing; p=0.077, Mann-Whitney test (n=45-47 images from N=2 transfections). **(E)**: Western blot of Caco-2 cells transfected with either control siRNA or NUDT1 siRNA. Blots were simultaneously probed with anti-ß-actin and anti-NUDT1 antibodies. Arrows mark the expected molecular weight of ß-actin or NUDT1. Right: Quantification of NUDT1 band intensity normalized to the ß-actin band intensity *: p<0.05, t-test (n=6-6 samples from N=2 transfections). **(F)**: Caco-2 monolayers were transfected with either non-targeting control siRNA or siRNA targeting NUDT1. Cells were fixed and used for anti-NUDT1 immunostaining 48 hours after transfection. Left: Representative images of anti-NUDT1 immunostaining. Right: quantification of NUDT1immunostaining. *: p<0.05, Student’s t-test (n=62-65 images from N=2 transfections).

**Figure S4.**
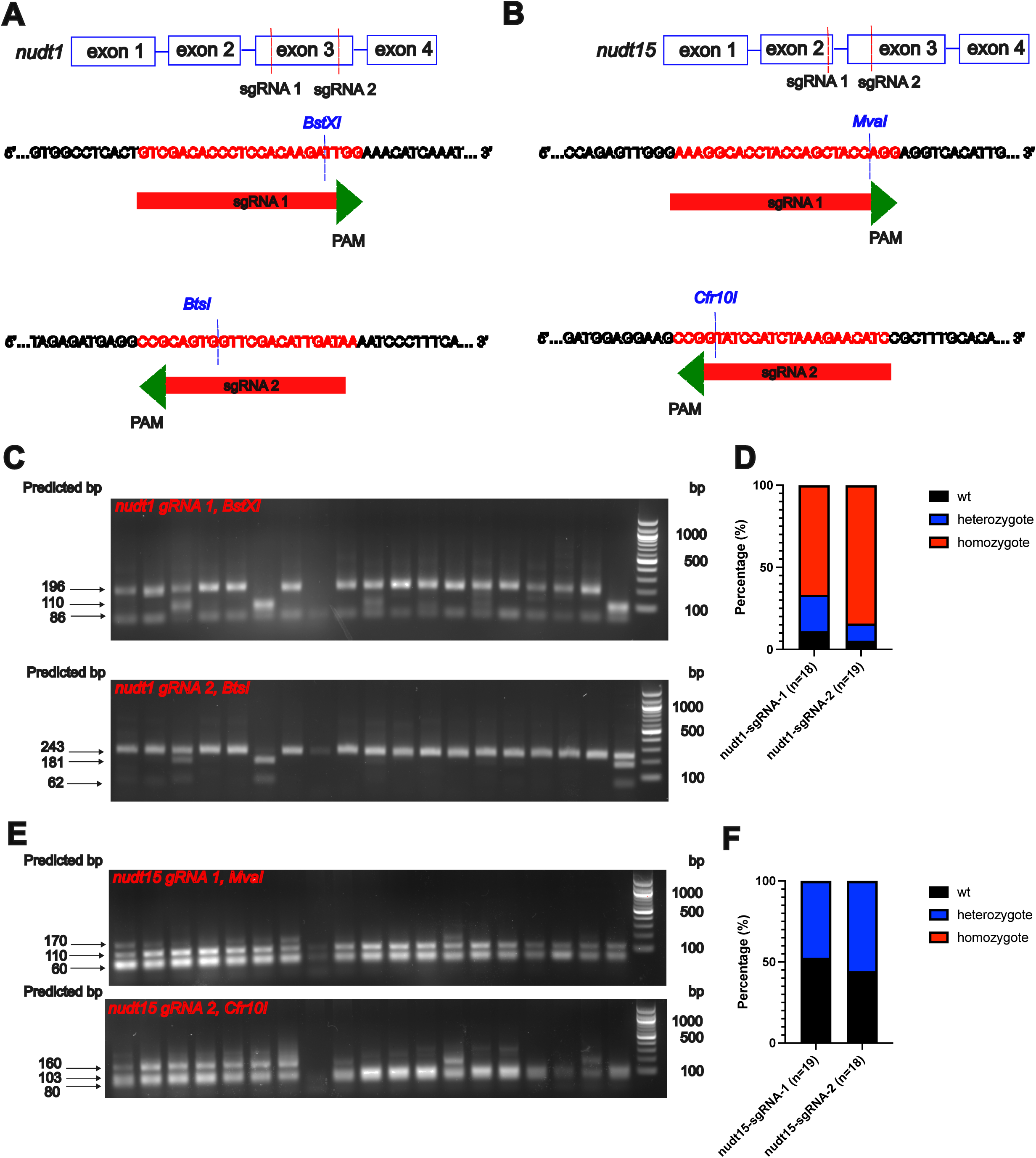
Extended data supporting Figure 4. **(A, B)** Schematic representation of zebrafish *nudt1* and *nudt15*. 2 guide RNAs (red text) were designed to disrupt exon2 (*nudt1)* and exons 2 and 3 (*nudt15*) and to disrupt restriction enzyme recognition sites (cleavage site marked with a blue dash) in these genes. **(C, E)** *nudt1* and *nudt15* F0 CRISPR larvae (18 or 19 per mutant) were genotyped by PCR amplification, restriction analysis, and agarose gel electrophoresis. For *nudt1* guide RNA-1, the digest of the wild type samples yields two lower bands of 110 and 86bp, digested heterozygotes have an additional higher band of 196 bp and homozygotes are uncleaved and result in only the 196bp band. For *nudt1* guide RNA-2, the digest of wild type samples results in two bands of 181 and 62 bp, heterozygotes have an additional higher band of 243 bp, while PCR products from homozygotes are uncleaved and only have the 243 bp band. For *nudt15* guide RNA-1, digested PCR products from wild type samples yield two bands of 100 and 60 bp, heterozygotes have an additional 170bp band, while in the case of the homozygotes, only the 170bp band is visible. For *nudt15* guide RNA-2, digested PCR products from wild type samples yield two bands of 103 and 80 bp (with an additional 57 bp fragment that can’t be visualized properly), heterozygotes have an additional 160 bp band, while in the case of the homozygotes, only the 160bp and 80bp band is visible. **(D, F)** The fraction of F0 CRISPR homozygotes, heterozygotes, or wt animals among the 18 or 19 tested larvae in C and D.

**Figure S5.**
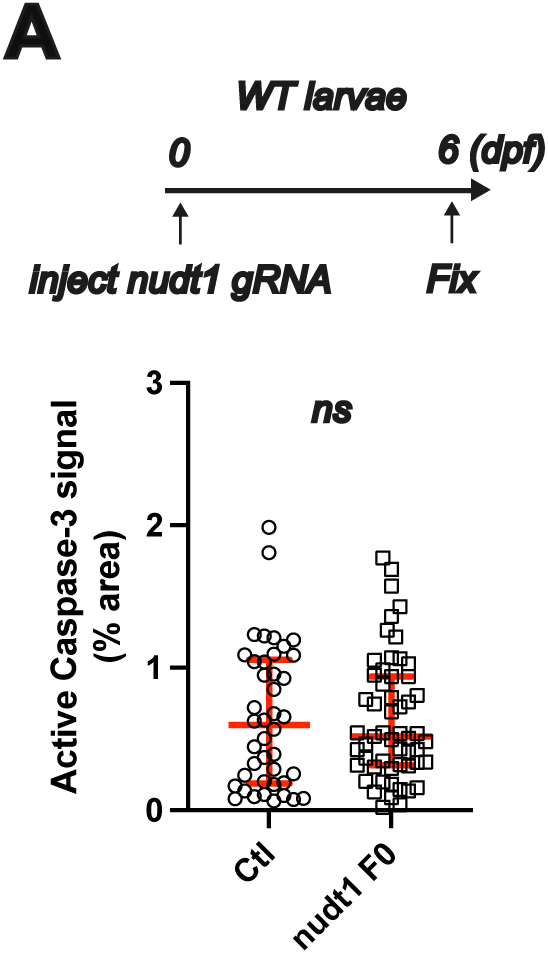
Extended data supporting Figure 4. **(A)** Top: Experimental scheme. WT zebrafish embryos were injected with Cas9 and guide RNAs *for nudt1* at the one cell stage. Larvae were fixed at 6dpf and used for anti-active caspase-3 immunostaining. Bottom: Quantification of apoptosis in control and F0 injected larvae. *Nudt1* F0 CRISPR had no effect on intestinal epithelial apoptosis; p=0.85, Mann-Whitney test (n= 42 and 55 larvae).

## References

1. Dupre-Crochet, S., M. Erard, and O. Nubetae, ROS production in phagocytes: why, when, and where? J Leukoc Biol, 2013. 94(4): p. 657–70.

2. Weavers, H. and P. Martin, The cell biology of inflammation: From common traits to remarkable immunological adaptations. J Cell Biol, 2020. 219(7).

3. Pena, O.A. and P. Martin, Cellular and molecular mechanisms of skin wound healing. Nat Rev Mol Cell Biol, 2024. 25(8): p. 599–616.

4. Sies, H. and D.P. Jones, Reactive oxygen species (ROS) as pleiotropic physiological signalling agents. Nat Rev Mol Cell Biol, 2020. 21(7): p. 363–383.

5. de Almeida, A., et al., ROS: Basic Concepts, Sources, Cellular Signaling, and its Implications in Aging Pathways. Oxid Med Cell Longev, 2022. 2022: p. 1225578.

6. Wang, Y., J. Yang, and J. Yi, Redox sensing by proteins: oxidative modifications on cysteines and the consequent events. Antioxid Redox Signal, 2012. 16(7): p. 649–57.

7. Erlemann, K.R., et al., Airway epithelial cells synthesize the lipid mediator 5-oxo-ETE in response to oxidative stress. Free Radic Biol Med, 2007. 42(5): p. 654–64.

8. Grant, G.E., et al., 5-oxo-ETE is a major oxidative stress-induced arachidonate metabolite in B lymphocytes. Free Radic Biol Med, 2011. 50(10): p. 1297–304.

9. Grant, G.E., et al., Enhanced formation of 5-oxo-6,8,11,14-eicosatetraenoic acid by cancer cells in response to oxidative stress, docosahexaenoic acid and neutrophil-derived 5-hydroxy-6,8,11,14-eicosatetraenoic acid. Carcinogenesis, 2011. 32(6): p. 822–8.

10. Kowal, K., A. Gielicz, and M. Sanak, The effect of allergen-induced bronchoconstriction on concentration of 5-oxo-ETE in exhaled breath condensate of house dust mite-allergic patients. Clin Exp Allergy, 2017. 47(10): p. 1253–1262.

11. Bowers, R., et al., Oxidative Stress in Severe Pulmonary Hypertension. American Journal of Respiratory and Critical Care Medicine, 2004. 169(6): p. 764–769.

12. Balgoma, D., et al., Linoleic acid-derived lipid mediators increase in a female-dominated subphenotype of COPD. Eur Respir J, 2016. 47(6): p. 1645–56.

13. Ma, Y., et al., Oxoeicosanoid signaling mediates early antimicrobial defense in zebrafish. Cell Rep, 2023. 42(1): p. 111974.

14. Enyedi, B., et al., Tissue damage detection by osmotic surveillance. Nat Cell Biol, 2013. 15(9): p. 1123–30.

15. Oehlers, S.H., et al., Retinoic acid suppresses intestinal mucus production and exacerbates experimental enterocolitis. Disease Models \& Mechanisms, 2012. 5(4): p. 457–467.

16. Oehlers, S.H., et al., Chemically induced intestinal damage models in zebrafish larvae. Zebrafish, 2013. 10(2): p. 184–193.

17. Katikaneni, A., et al., Lipid peroxidation regulates long-range wound detection through 5-lipoxygenase in zebrafish. Nat Cell Biol, 2020. 22(9): p. 1049–1055.

18. Sharrock, A.V., et al., NTR 2.0: a rationally engineered prodrug-converting enzyme with substantially enhanced efficacy for targeted cell ablation. Nat Methods, 2022. 19(2): p. 205–215.

19. Hashiguchi, K., et al., The roles of human MTH1, MTH2 and MTH3 proteins in maintaining genome stability under oxidative stress. Mutat Res, 2018. 808: p. 10–19.

20. Sakumi, K., et al., Cloning and expression of cDNA for a human enzyme that hydrolyzes 8-oxo-dGTP, a mutagenic substrate for DNA synthesis. Journal of Biological Chemistry, 1993. 268(31): p. 23524–23530.

21. Pochard, C., et al., PGI2 inhibits intestinal epithelial permeability and apoptosis to alleviate colitis. Cellular and Molecular Gastroenterology and Hepatology, 2021. 12(3): p. 1037–1060.

22. Gao, J., et al., Integrative analysis of complex cancer genomics and clinical profiles using the cBioPortal. Sci Signal, 2013. 6(269): p. pl1.

23. White, R.M., et al., Transparent adult zebrafish as a tool for in vivo transplantation analysis. Cell Stem Cell, 2008. 2(2): p. 183–9.

24. Arnaud, E., et al., The zebrafish bcl-2 homologue Nrz controls development during somitogenesis and gastrulation via apoptosis-dependent and -independent mechanisms. Cell Death Differ, 2006. 13(7): p. 1128–37.

25. Williams, J.A., et al., Programmed cell death in zebrafish rohon beard neurons is influenced by TrkC1/NT-3 signaling. Dev Biol, 2000. 226(2): p. 220–30.

26. Gelashvili, Z., et al., Perivascular Macrophages Convert Physical Wound Signals Into Rapid Vascular Responses. bioRxiv, 2024.

27. Riemslagh, F.W., et al., Reduction of oxidative stress suppresses poly-GR-mediated toxicity in zebrafish embryos. Dis Model Mech, 2021. 14(11).

28. Rosowski, E.E., et al., Macrophages inhibit Aspergillus fumigatus germination and neutrophil-mediated fungal killing. PLoS Pathog, 2018. 14(8): p. e1007229.

29. Wurster, S., et al., EGF-mediated suppression of cell extrusion during mucosal damage attenuates opportunistic fungal invasion. Cell Rep, 2021. 34(12): p. 108896.

30. Cocchiaro, J.L. and J.F. Rawls, Microgavage of zebrafish larvae. J Vis Exp, 2013(72): p. e4434.

31. Kwan, K.M., et al., The Tol2kit: a multisite gateway-based construction kit for Tol2 transposon transgenesis constructs. Dev Dyn, 2007. 236(11): p. 3088–99.

32. Tamas, S.X., et al., A genetically encoded sensor for visualizing leukotriene B4 gradients in vivo. Nat Commun, 2023. 14(1): p. 4610.

33. Murdoch, C.C., et al., Intestinal Serum amyloid A suppresses systemic neutrophil activation and bactericidal activity in response to microbiota colonization. PLoS Pathogens, 2019. 15(3): p. e1007381.

34. Carreras-Puigvert, J., et al., A comprehensive structural, biochemical and biological profiling of the human NUDIX hydrolase family. Nat Commun, 2017. 8(1): p. 1541.

35. Kalyvianaki, K., et al., Antagonizing effects of membrane-acting androgens on the eicosanoid receptor OXER1 in prostate cancer. Sci Rep, 2017. 7: p. 44418.

36. Yu, T., et al., Kruppel-like factor 4 regulates intestinal epithelial cell morphology and polarity. PLoS One, 2012. 7(2): p. e32492.

